# Effects of pro-depressant and immunomodulatory drugs on biases in decision-making in the rat judgement bias task

**DOI:** 10.1101/2020.10.28.358655

**Authors:** CA Hales, JM Bartlett, R Arban, B Hengerer, ESJ Robinson

## Abstract

Studies in human and non-human species suggest that decision-making behaviour can be biased by affective state, also termed an affective bias. To study these behaviours in non-human species, judgement bias tasks have been developed. Animals are trained to associate specific cues (tones) with a positive or negative/less positive outcome. Animals are then presented with intermediate ambiguous cues and affective biases quantified by observing whether animals make more optimistic or more pessimistic choices. Here we use a high versus low reward judgement bias task and test whether pharmacologically distinct compounds, which induce negative biases in learning and memory, have similar effects on decision-making: tetrabenazine (0.0-1.0mg/kg), retinoic acid (0.0-10.0mg/kg) and rimonabant (0.0-10.0mg/kg). We also tested immunomodulatory compounds: interferon-α (0-100units/kg), lipopolysaccharide (0.0-10.0μg/kg) and corticosterone (0.0-10.0mg/kg). We observed no specific effects in the judgement bias task with any acute treatment except corticosterone which induced a negative bias. We have previously observed a similar lack of effect with acute but not chronic psychosocial stress and so next tested decision-making behaviour following chronic interferon-alpha. Animals developed a negative bias which was sustained even after treatment was ended. These data suggest that decision-making behaviour in the task is sensitive to chronic but not acute effects of most pro-depressant drugs or immunomodulators, but exogenous administration of acute corticosterone induces pessimistic behaviour. This work supports our hypothesis that biases in decision-making develop over a different temporal scale to those seen with learning and memory which may be relevant in the development and perpetuation of mood disorders.

**Graphical abstract and text:** Decision-making bias in rats, measured using a judgement bias task, is not altered by acute treatments with pro-depressant or immunomodulatory drugs, but becomes more negative following chronic treatment. The time course of change in decision-making bias reflects the subjective reporting of changes in depression symptoms in humans treated with these drugs.

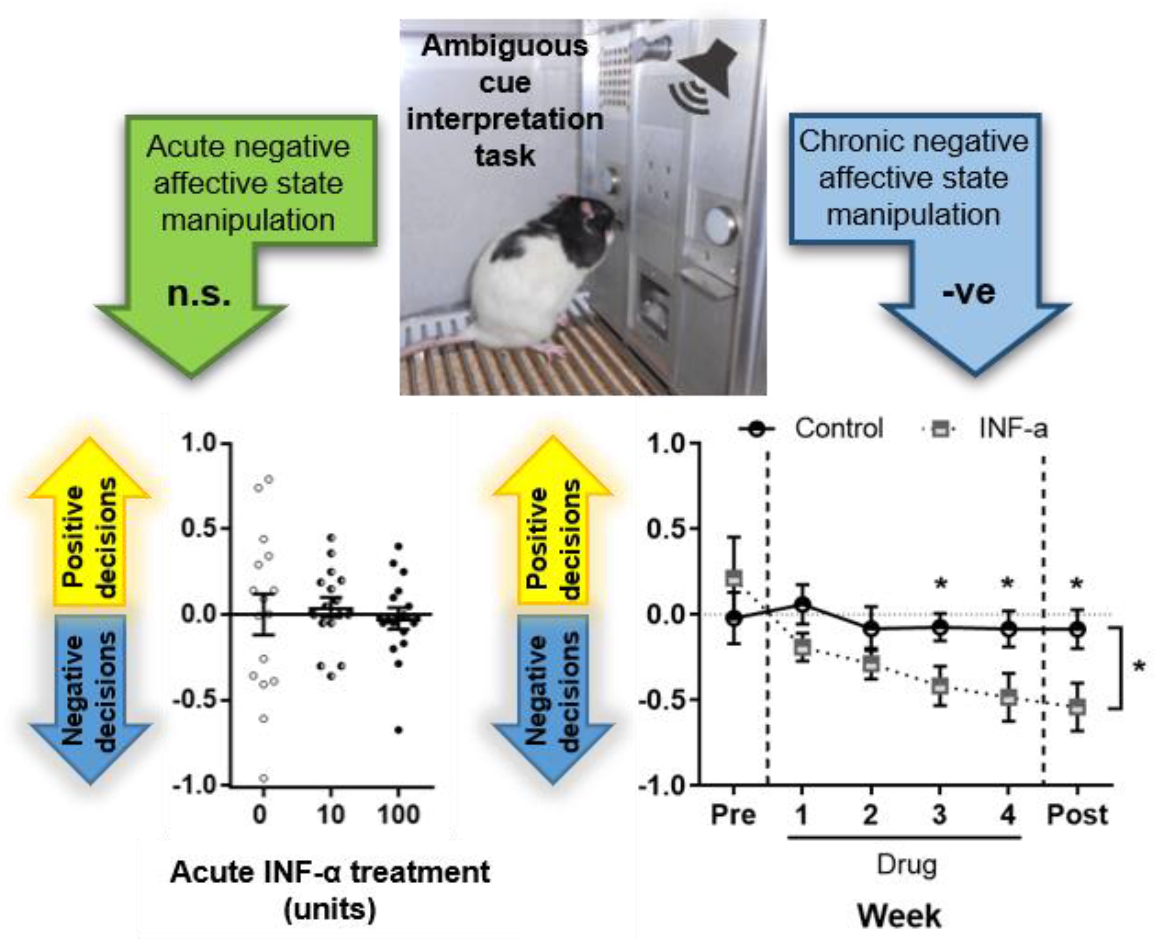

## Introduction

Affective biases, when emotions alter cognitive processing, occur across many different cognitive domains. Studies have demonstrated that negative affective biases in processes such as emotional interpretation, learning, memory and decision-making contribute to the development and maintenance of mood disorders such as depression and anxiety^1-5^. In healthy participants and in major depressive disorder (MDD), it has been shown that positive biases in emotional processing can be induced following acute treatments with antidepressants, despite a lack of subjectively reported change in mood^6^. These findings support earlier hypotheses relating to the role of neuropsychological processes in MDD^7,8^, and adds to the proposal that negative affective biases have a causal role in the development, maintenance and treatment of MDD^9-11^. This theory also posits that pharmacological treatments may work by remediating negative processing of information that, over time, leads to symptomatic improvements. Therefore, investigating the time courses and mechanisms that underlie changes in affective biases may provide further insight into the underlying psychology of mood disorders.

Affective biases can be measured in animal models^12-14^. The judgement bias task (JBT; also known as the ambiguous cue interpretation task) is a rodent decision-making task that measures biases in interpretation of ambiguous cues^12,15^. The task has been reverse-translated for use in humans and has shown translational validity^16-18^. In humans, self-reported state anxiety correlated with degree of negative bias^16^, and people with pathological anxiety symptoms exhibited the same negative biases in the task^17^ to those observed in rats that have experienced anxiogenic manipulations^19^. Furthermore, individual differences in decision-making bias have been shown to be reliably linked to individual differences in depression symptoms^18^. Studies in putative models of depression suggest that rats in negative affective states make more pessimistic choices during ambiguous cue presentation^19-23^. Studies with pharmacological treatments, including antidepressant drugs have resulted in a more mixed picture^24^. Our work in rats has revealed differences between the time course of biases seen following treatment with conventional antidepressants compared rapid acting antidepressants (RAAD) that have shown efficacy in clinical settings, including ketamine^25,26^, an NMDA receptor antagonist; CP-101,606, a GluN2B receptor subunit antagonist; and scopolamine, a muscarinic receptor antagonist^26^. When given acutely fluoxetine, reboxetine and venlafaxine (conventional antidepressants) had no effect on bias, but chronic treatment with fluoxetine resulted in a positive judgement bias that was seen in the second and third weeks of treatment^26^. This contrasts with acute ketamine, CP-101,606 or scopolamine treatment, which all induce an immediate positive judgement bias^25,26^. Other NMDA receptor antagonists that have not shown clinical antidepressant efficacy (PCP, lanicemine and memantine) also fail to induce a change in bias when given acutely^25,26^. These data suggest that this reward-based JBT is sensitive to pharmacological treatments that induce biases across time courses that correspond to subjectively reported change in mood in humans following these drug treatments.

In contrast, another rodent task that measures affective biases in learning and memory, the affective bias test (ABT)^27^, is sensitive to acute changes in affective state induced by conventional antidepressants, as in humans^6^. In the ABT a dissociation between conventional antidepressants and RAADs is also observed^28^. Acute treatments with conventional antidepressants positively biased new learning but failed to attenuate previously acquired negative biases when administered immediately before testing memory recall^27,28^, whereas ketamine had the opposite effect^28^. It has also been shown that acute treatment with putative pro-depressant treatments, including drugs with distinct pharmacological mechanisms that have been linked to increased risk of depression in the clinic, and immunomodulators that alter immune system function, are able to induce negative biases in this task^27,29^.

In this study we investigate whether the putative pro-depressant drugs that induce negative biases in learning and memory when given acutely in the ABT have the same effect on decision-making biases in the JBT. Specifically, we tested rimonabant, the anti-obesity drug that was withdrawn from the market following evidence that it causes an increased risk of suicidal tendencies and depression^30^; retinoic acid, the active ingredient of the acne drug Roaccutane that has been associated with an increased incidence of depression in patients^31^; and tetrabenazine, a vesicular monoamine transport inhibitor used as an off-label treatment for chorea in Huntington’s disease and has also been associated with adverse psychiatric symptoms^32,33^. We also tested the immunomodulators interferon-α (IFN-α), an immunotherapy drug shown to increase the risk of depression and suicidality in patients^34,35^; lipopolysaccharide (LPS), the proinflammatory mediator used chronically depression model in rodents^36^; and corticosterone, the rodent stress hormone that has also been shown to induce depression-like behaviour in rodents following chronic treatment^37^. In previous studies using psychosocial stress we observed negative decision-making biases in the JBT following chronic but not acute exposure^19^, hence here we also tested interferon-α effects following chronic treatment.

## Methods

### Animals and apparatus

Three cohorts of male Lister Hooded rats (cohort 1: n=16; cohort 2: n=16; cohort 3: n=16) were used (Harlan, UK). Rats weighed 260-305g (cohort 1) / 270-305g (cohort 2) / 275-295g (cohort 3) at the start of training, and 305-445g (cohort 1) / 400-465g (cohort 2) / 390-460g (cohort 3) by the start of experimental manipulations. Rats were housed in pairs with environmental enrichment consisting of a red Perspex house, cardboard tube, wood chew block and rope tied across the cage lid. Rats were kept under temperature (19-23°C) and humidity (45-65%) controlled conditions on a 12h reverse lighting cycle (lights off at 08:00h). Water was available *ad libitum* in the home cage, but rats were maintained at no less than 90% of their free-feeding body weight, matched to a standard growth curve, by restricting access to laboratory chow (LabDiet, PMI Nutrition International) to ∼18g per rat per day. All procedures were carried out under local institutional guidelines (University of Bristol Animal Welfare and Ethical Review Board) and in accordance with the UK Animals (Scientific Procedures) Act 1986. During experiments all efforts were made to minimise suffering, and at the end of experiments rats were killed by giving an overdose of sodium pentobarbitone (200mg/kg). Behavioural testing was carried out between 0800-1800h, using standard rat operant chambers (Med Associates, Sandown Scientific, UK) as previously described^19,25,26^. Operant chambers (30.5×24.1×21.0cm) used for behavioural testing were housed inside a light-resistant and sound-attenuating box. They were equipped with two retractable response levers positioned on each side of the centrally located food magazine. The magazine had a house light (28V, 100mA) located above it. An audio generator (ANL-926, Med Associates, Sandown Scientific, UK) produced tones that were delivered to each chamber via a speaker positioned above the left lever. Operant chambers and audio generators were controlled using K-Limbic software (Conclusive Solutions Ltd., UK).

### Behavioural task

Animals were tested using a high versus low reward version of the JBT as previously reported^19,25,26^. Rats were first trained to associate one tone (2kHz at 83dB rats, designated high reward) with a high value reward (four 45mg reward pellets; TestDiet, Sandown Scientific, UK) and the other tone (8kHz at 66dB, designated low reward) with a low value reward (one 45mg reward pellet) if they pressed the associated lever (either left or right, counterbalanced across rats) during the 20s tone (see Figure S1 for a detailed depiction of the task). Response levers were extended at the beginning of every session and remained extended for the duration of the session (maximum one hour for all session types). All trials were self-initiated via a head entry into the magazine, followed by an intertrial interval (ITI), and then presentation of the tone. Pressing the incorrect lever during a tone was punished by a 10s timeout, as was an omission if the rat failed to press any lever during the 20s tone. Lever presses during the ITI (prematures) were punished by a 10s timeout. During a timeout, the house light was illuminated, and responses made on levers were recorded but had no programmed consequences.

### Training

Training was the same for all cohorts. Table S1 contains a summary of training stages used, but briefly were as follows:

1. Magazine training: tone played for 20s followed by release of one pellet into magazine. Criteria: 20 pellets eaten for each tone frequency.
2. Tone training: response on lever during tone rewarded with one pellet. Only one tone frequency, and one lever available per session. Criteria: >50 trials completed.
3. Discrimination training: response on correct corresponding lever only during tone rewarded with one pellet. Both tones played (pseudorandomly) and both levers available. Criteria: >70% accuracy for both tones, <1:1 ratio of correct:premature responses and no significant difference on any behavioural measures analysed over three sessions.
4. Reward magnitude training: As for discrimination training but 2kHz tone now rewarded with four pellets, 8kHz tone rewarded with one pellet. Criteria: as for discrimination training but with >60% accuracy for both tones.

Rats were required to meet criteria for at least two consecutive sessions before progressing to the next training stage. Once trained (29 total sessions, see Table S1 for number of sessions required for each training stage), animals were used in judgement bias experiments.

### Judgement bias testing

Baseline sessions (100 trials: 50 high and 50 low reward tones; pseudorandomly, for details see Table S1) were conducted on Monday and Thursday. Probe test sessions (120 trials: 40 high reward, 40 low reward, and 40 ambiguous midpoint tones; pseudorandomly, for details see Table S1) were conducted on Tuesday and Friday. The midpoint tone was randomly reinforced whereby 50% of trials had outcomes as for the high reward tone, and 50% as for the low reward tone. This was to ensure a specific outcome could not be learnt, and to maintain responding throughout the experiments (see Figure S1 and Table S1 for a detailed description of how this was implemented). All rats were initially trained and tested using a 5kHz (75dB) midpoint tone. Cohort 1 (n=16) were then used to test the acute effect of treatment with corticosterone (Table 1). Following this, half of these rats (cohort 1a, n=8; Table 1) were used for another experiment, while the other half (cohort 1b, n=8) were then used for the remaining acute treatments conducted in this study (listed in Table 1). After being split, it was found that cohort 1b displayed more negative baseline interpretation of the midpoint ambiguous tone (see Figure S2). As drugs hypothesised to induce negative affective state were to be tested in these rats, these animals were subsequently switched to a 4.75 kHz ambiguous midpoint tone to prevent a “floor effect” for the remaining manipulations (i.e. to allow room for drugs to cause more negative responding; Figure S2). Rats in cohort 2 were initially used to test the effect of acute treatments with NMDA receptor antagonists on judgement bias of the 5kHz midpoint tone (experiments not reported here; see Hales et al.^26^). These rats then went on to be used to test the effects of acute treatments listed in Table 1 along with cohort 1b, and so cohort 2 were also moved to a 4.75kHz ambiguous midpoint tone to match (Figure S2). Cohort 3 were used to test the effect of chronic treatment with INF-α.

**Table 1.** Drug treatments given to each cohort of rats.

| Cohort | # rats | Midpoint tone frequency | Treatment | Doses (mg/kg unless otherwise stated) |
| --- | --- | --- | --- | --- |
| **1** | 16 | 5 kHz | Corticosterone | 0, 1, 10 |
| **1a** | 8 | 5 kHz | Used in a chronic study (not reported here) |  |
| **1b** | 8 | 4.75 kHz | Lipopolysaccharide<br>Retinoic acid<br>Rimonabant<br>Tetrabenazine | 0, 1, 3, 10 μg/kg<br>0, 3, 10<br>0, 3, 10<br>0.0, 0.3, 1.0 |
| **2** | 16 | 4.75 kHz | Interferon-α (acute)<br>Lipopolysaccharide<br>Retinoic acid<br>Rimonabant<br>Tetrabenazine | 0, 10, 100 units/kg<br>0, 1, 3, 10 μg/kg<br>0, 3, 10<br>0, 3, 10<br>0.0, 0.3, 1.0 |
| **3** | 16 | 5 kHz | Chronic interferon-α | 0, 100 units/kg (daily) |

### Experimental design and drugs

All acute dose-response studies used a within-subject fully counterbalanced drug treatment schedule (see Table 1 for details of individual treatments). All drugs (except corticosterone) were given by intraperitoneal injection using a low-stress, non-restrained technique^38^. Corticosterone (Sigma Aldrich, UK) was dissolved in 5% DMSO and 95% sesame oil and administered by subcutaneous injection 30 minutes prior to testing. Rimonabant (kindly provided by Pfizer) and 13-cis retinoic acid (Sigma Aldrich, UK) were dissolved in 5% DMSO, 10% cremaphor and 85% sterile saline and given 30 and 60 minutes (respectively) prior to testing. Tetrabenazine was dissolved in 20% DMSO and 80% saline at pH 2.0 which was then adjusted to pH 5.5 for dosing, and given 30 minutes prior to testing. IFN-α and LPS were resuspended in saline and stocks stored at -20°C until use. These were also given 30 minutes prior to testing. Drug doses were selected based on previous rodent behavioural studies, particularly the ABT at doses that had shown efficacy^27,29^. For all studies, the experimenter was blind to drug dose. Each dose-response study was separated by at least one week (five sessions) of baseline testing.

For the chronic INF-α experiment, a between-subjects study design was used. This was split into three parts: (1) a pre-drug week, (2) four weeks of drug treatment, and (3) one week post-drug testing. Rats were split into control (0.9% sterile saline vehicle) or INF-α (100units/kg) groups based on task performance (matched for all analysed behavioural variables) during the pre-drug week. The experimenter was blind to treatment group. Rats were dosed daily 30 minutes prior to behavioural testing (or at an equivalent time on days when behavioural testing did not occur) by intraperitoneal injection using a low-stress, non-restrained technique^38^. Treatment commenced the Monday of first drug week and ended on the Friday of the final drug week.

### Data and statistical analysis

Sample size was estimated based on our previous studies using the JBT^19,25,26^. Changes in bias should occur without effects on other variables, therefore strict inclusion criteria were established to reduce any potential confound in the data analysis. Only animals which maintained more than 60% accuracy for each reference tone, less than 50% omissions, and also completed more than 50% of the total trials were used for analysis. Details of animals excluded from each study are given in Table 2. Cognitive bias index (CBI) was used as a measure of judgement bias in response to the midpoint tone. CBI was calculated by subtracting the proportion of responses made on the low reward lever from the proportion made on the high reward lever. This created a score between -1 and 1, where negative values represent a negative bias and positive values a positive bias. Change from baseline in CBI was then calculated for all experimental manipulations as follows: vehicle (0.0mg/kg) probe test CBI − drug dose probe test CBI, for acute experiments; and pre-drug week probe test CBI – drug week probe test CBI for the chronic study. This was calculated to take into account individual differences in baseline bias, and to make directional changes caused by drug treatments clearer. Although individuals within a cohort were variable regarding their CBI scores at baseline (see Figure S3 for raw CBI data for all acute drug studies), performance was consistent across repeated sessions. To provide individual values for vehicle probe test sessions for this measure, the population average for this session was taken away from each individual rats’ CBI score for the same session. This allowed analysis with repeated measures analysis of variance (rmANOVA) with session as the within-subjects factor for acute studies, and a mixed ANOVA with the addition of group as the between-subjects factor for the chronic study.

Response latency and percentages of positive responses, omissions and premature responses were also analysed (see Table S2 for details). For acute drug studies, these were analysed with rmANOVAs with session and tone as the within-subjects factors. The chronic study was analysed similarly, but with the addition of group as a between-subjects factor. Paired t-tests (acute studies) and/or independent samples t-tests (chronic study) were performed as post-hoc tests if significant effects were established. Huynh-Feldt corrections were used to adjust for violations of the sphericity assumption, and Sidak correction was applied for multiple comparisons. All statistical tests were conducted using SPSS 24.0.0.2 for Windows (IBM SPSS Statistics) with α=0.05. Results are reported with the ANOVA F-value (degrees of freedom, error) and *p*-value as well as any post-hoc *p*-values. All graphs were made using Graphpad Prism 7.04 for Windows (Graphpad Software, USA).

**Table 2.** Details of rats excluded for each drug treatment.

| Drug | Cohort(s) | Total number rats | Rats excluded from analysis | N number for final analysis |
| --- | --- | --- | --- | --- |
| **Tetrabenazine** | 1b and 2 | 24 | 7: failure to complete sufficient trials on the 1 mg/kg dose | 17 |
| **Rimonabant** | 1b and 2 | 24 | 3: failure to complete sufficient trials on the 3 mg/kg dose<br>(10 mg/kg dose completely excluded as the majority of animals failed to complete sufficient trials) | 21 |
| **Retinoic acid** | 1b and 2 | 24 | 0 | 24 |
| **Interferon-α** | 2 | 16 | 0 | 16 |
| **LPS** | 1b and 2 | 24 | 1: did not meet accuracy criteria on vehicle session | 23 |
| **Corticosterone** | 1 | 16 | 0 | 16 |
| **Chronic interferon-α** | 3 | 16 (15) | 15 rats began the drug study as one animal died from an unrelated health issue during training. 1 rat was excluded from the INF-α group for failing to meet accuracy criteria and 1 animal was euthanised during the study as it developed seizures | 13 (n = 8 control, n = 5 INF-α) |

## Results

### Effects of acute treatment with putative pro-depressant drugs on interpretation of the ambiguous cue in the JBT

For tetrabenazine, seven rats were excluded from the final analysis as they failed to complete sufficient trials on the 1.0mg/kg dose. For rimonabant, the highest dose (10.0mg/kg) had to be excluded from the final analysis as only four rats completed sufficient trials. Three rats were excluded from the rest of the data analysis because they failed to complete sufficient trials on 3.0mg/kg dose. No rats were excluded from analysis for retinoic acid. The full datasets for behavioural measures without these exclusions can be found in Figures S4-5.

Tetrabenazine, rimonabant and retinoic acid did not change CBI at any of the doses tested (Figures 1-3, panel A). For tetrabenazine, there was a main effect of session for response latency (*F*_2,32_=3.593, *p*=0.039). Post-hoc analyses revealed a main effect of session for the high tone only (*F*_2,32_=3.508, *p*=0.042), with a trend towards a difference between the 0.0 and 1.0mg/kg doses (*p*=0.072; Figure 1B). There were no effects on other behavioural measures (Figure 1C-E). Rimonabant (3.0mg/kg) caused changes in responding to the reference tones (session*tone interaction: *F*_2,40_=4.240, *p*=0.021), whereby rats became less accurate for the high tone (*p*=0.045; Figure 2C), with a tendency for responding with greater accuracy for the low tone (*p*=0.062; Figure 2C). Rimonabant (3.0mg/kg) also increased omissions (main effect of session: *F*_1,20_=10.671, *p*=0.004 and session*tone interaction: *F*_1.515,_30.297=4.316, *p*=0.032) for the midpoint (*p*=0.002) and low (*p*=0.021) tones (Figure 2D). There was no effect on response latencies (Figure 2B) or premature responses (Figure 2E). Retinoic acid had no effect on any behavioural measures (Figure 3B-E).

**Figure 1.**
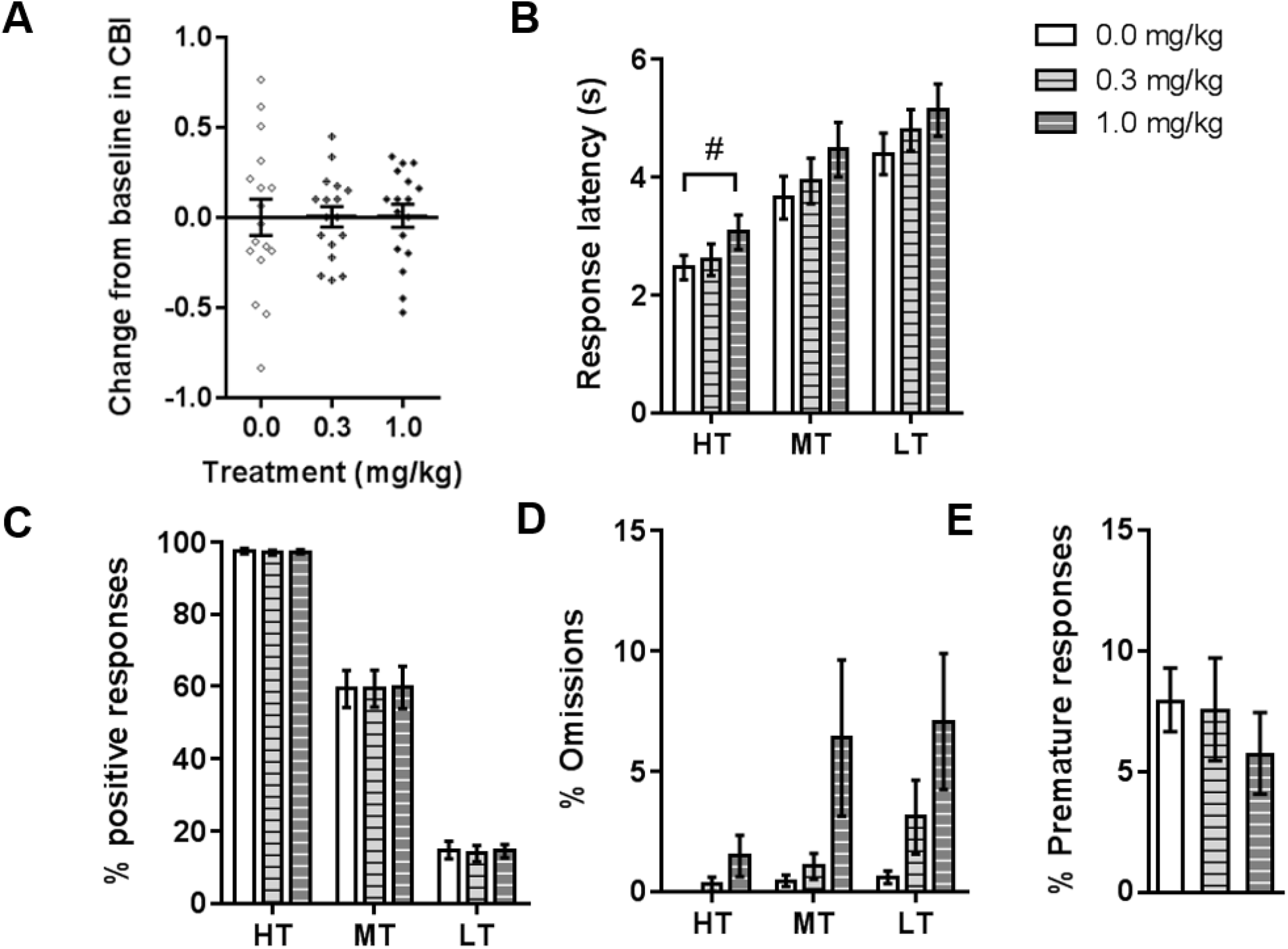
The effect of acute treatment with the putative pro-depressant drug tetrabenazine on judgement bias. Acute doses of tetrabenazine (0.0, 0.3, 1.0 mg/kg; n = 17) were administered by intraperitoneal injection prior to testing on the judgement bias task. (A) None of the doses tested induced a change in cognitive bias index (CBI) for the midpoint tone. (B) The higher dose of tetrabenazine (1.0 mg/kg) tended towards increasing response latency for the high reward tone (main effect of drug: *F*_2,32_=3.593, *p*=0.039 and post-hoc test: *p*=0.072). (B-D) There was no other effect of the doses tested on any other behavioural measures. Data shown and represent mean ± SEM (bars and error bars) overlaid with individual data points for each rat on panel A. #*p*<0.08. 30 min pre-treatment. HT - high reward tone; MT - midpoint tone; LT - low reward tone.

**Figure 2.**
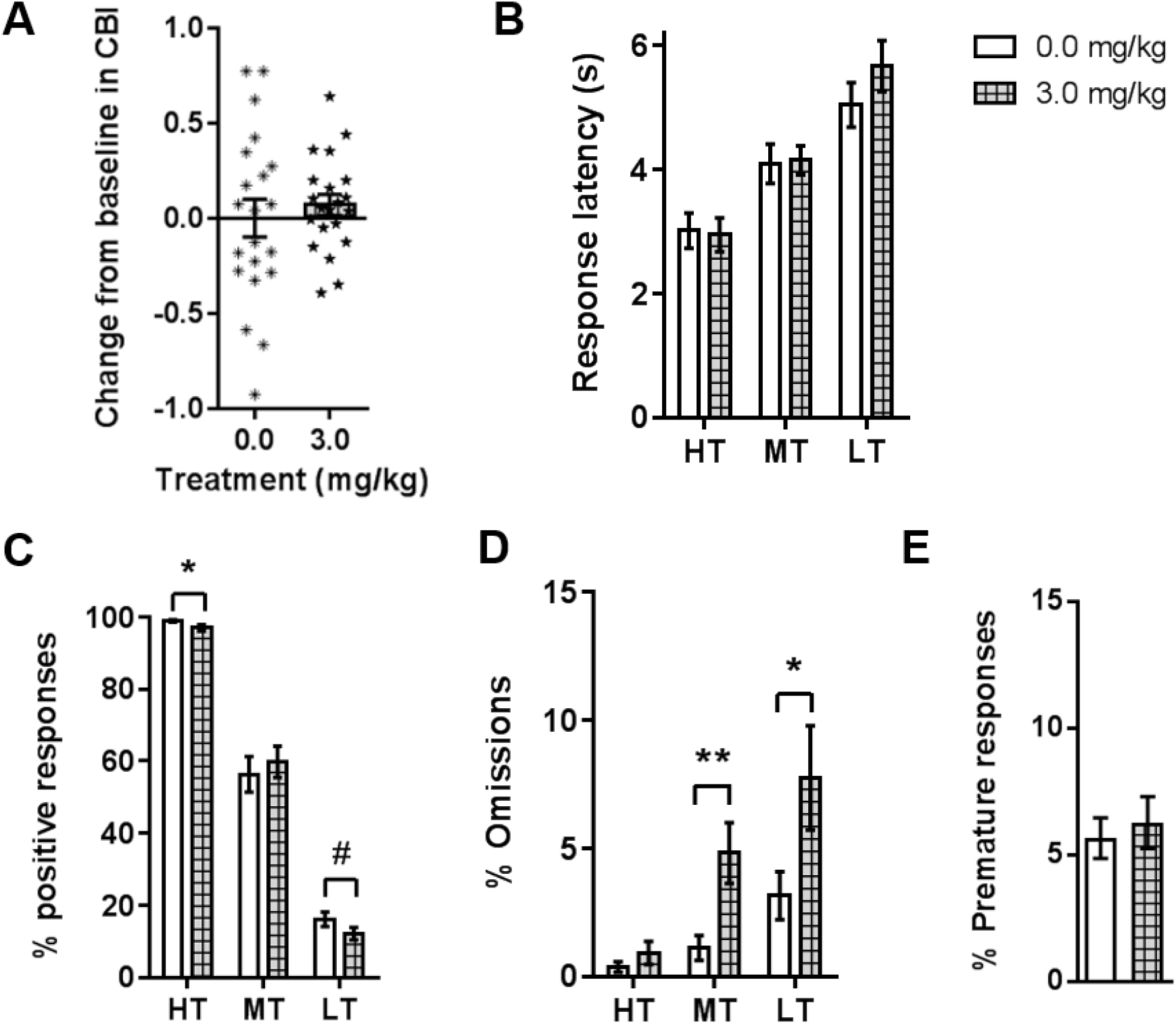
The effect of acute treatment with the putative pro-depressant drug rimonabant on judgement bias. Acute doses of rimonabant (0.0, 3.0 mg/kg; n = 21) were administered by intraperitoneal injection prior to testing on the judgement bias task. (A) Rimonabant induced a change in cognitive bias index (CBI) for the midpoint tone. (B, E) Rimonabant (3.0 mg/kg) had no effect on response latency or premature responding, but did cause (C) changes in positive responding (significant drug*tone interaction: *F*_2,40_=4.240, *p*=0.021). Accuracy was reduced for the high reward tone (*p*=0.045) whilst there was a tendency for accuracy to be increased for the low reward tone (*p*=0.062). (D) Rimonabant also increased omissions (significant drug*tone interaction: *F*_1.515,30.297_=4.316, *p*=0.032 for the midpoint (*p*=0.002) and low reward tones (*p*=0.021). Data shown and represent mean ± SEM (bars and error bars) overlaid with individual data points for each rat in panel A. \*\**p*<0.01; \**p*<0.05, #*p*<0.08. 30 min pre-treatment. HT - high reward tone; MT - midpoint tone; LT -low reward tone.

**Figure 3.**
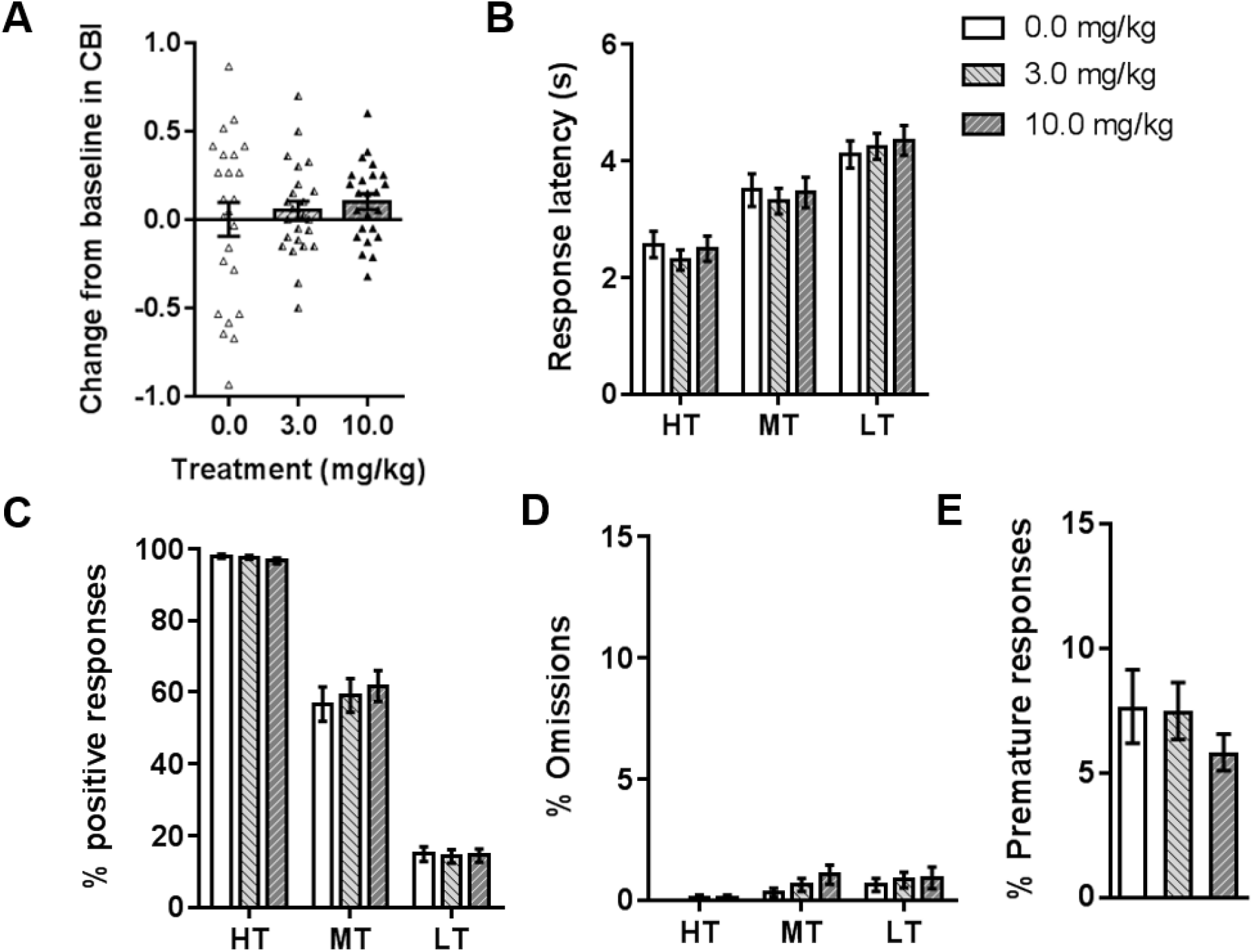
The effect of acute treatment with the putative pro-depressant drug retinoic acid on judgement bias. Acute doses of retinoic acid (0.0, 3.0, 10.0 mg/kg; n = 24) were administered by intraperitoneal injection prior to testing on the judgement bias task. (A-E) There were no effects of retinoic acid on any behavioural measures analysed. Data shown and represent mean ± SEM (bars and error bars) overlaid with individual data points for each rat on panel A. 60 min pre-treatment. HT - high reward tone; MT - midpoint tone; LT - low reward tone.

### Effects of acute treatment with immunomodulators on interpretation of the ambiguous cue in the JBT

One rat had to be excluded from the LPS drug study as it did not meet accuracy criteria for the reference tones on the 0.0mg/kg session. All rats were included in the analysis for the INF-α and corticosterone drug studies.

None of the doses of INF-α or LPS tested caused a change in CBI (Figure 4A and 5A). These drugs also did not alter any other behavioural measures (Figure 4B-E and 5B-E). Corticosterone (10.0mg/kg) caused a negative change in CBI for the midpoint tone (main effect of session: *F*_2,30_=4.493, *p*=0.020; post-hoc: *p*=0.030; Figure 6A). Corticosterone had no effect on other behavioural measures (Figure 6B/D/E), apart from the 10mg/kg dose reducing percentage positive responses for the midpoint tone (session*tone interaction: *F*_4,60_=2.612, *p*=0.044; post-hoc main effect of session for midpoint tone: *F*_2,30_=4.009, *p*=0.029; post-hoc comparison: *p*=0.026; Figure 6C), which reflects the effect seen on CBI (Figure 6A).

**Figure 4.**
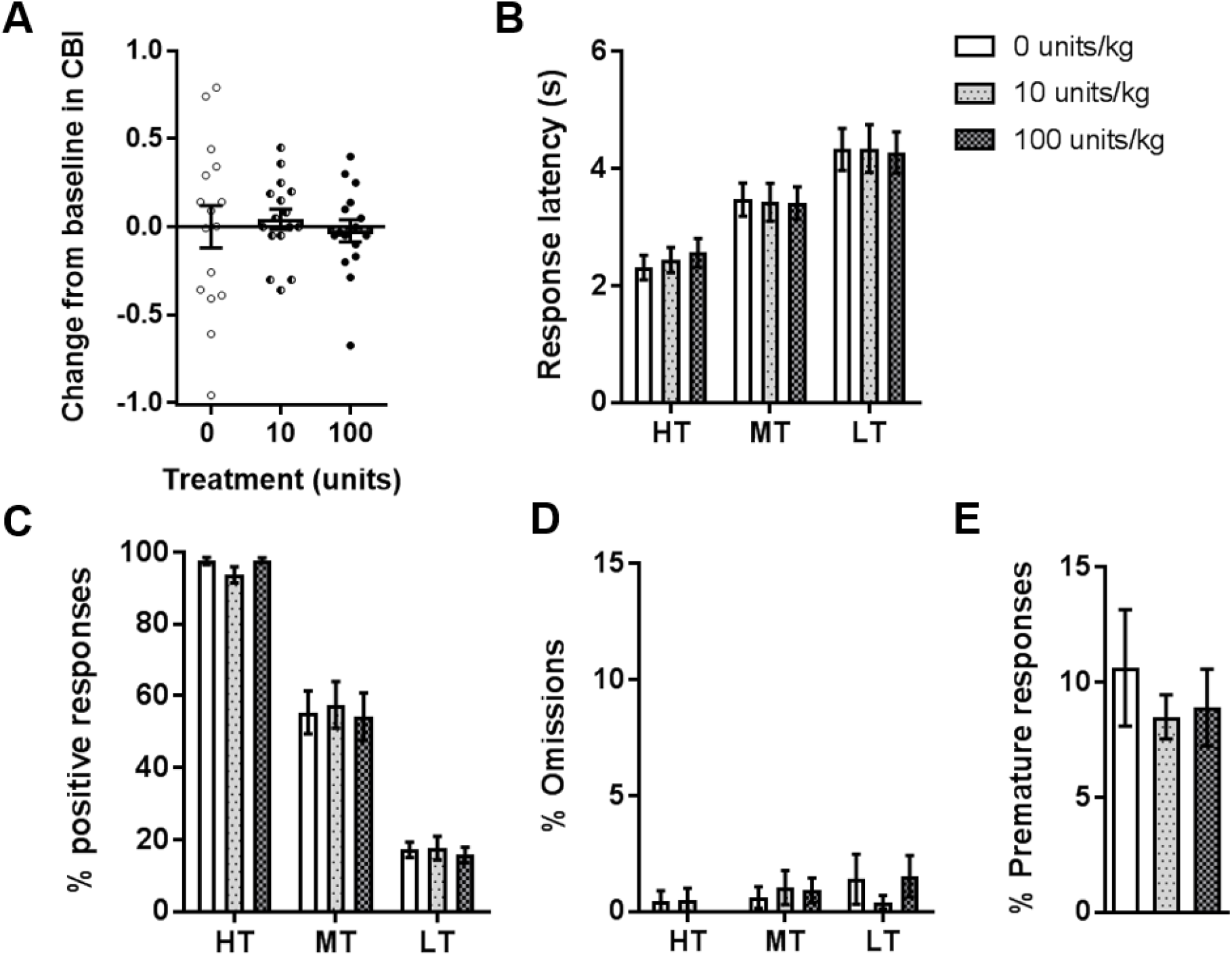
The effect of acute treatment with the immunomodulator interferon-α on judgement bias. Acute doses of interferon-α (IFN-α; 0, 10, 100 units/kg; n = 16) were administered by intraperitoneal injection prior to testing on the judgement bias task. (A) IFN-α did not induce a change in cognitive bias index (CBI) for the midpoint tone at the doses tested. (B-E) There was also no effect on other behavioural measures. Data shown and represent mean ± SEM (bars and error bars) overlaid with individual data points for each rat in panel A. 30 min pre-treatment. HT - high reward tone; MT - midpoint tone; LT - low reward tone.

**Figure 5.**
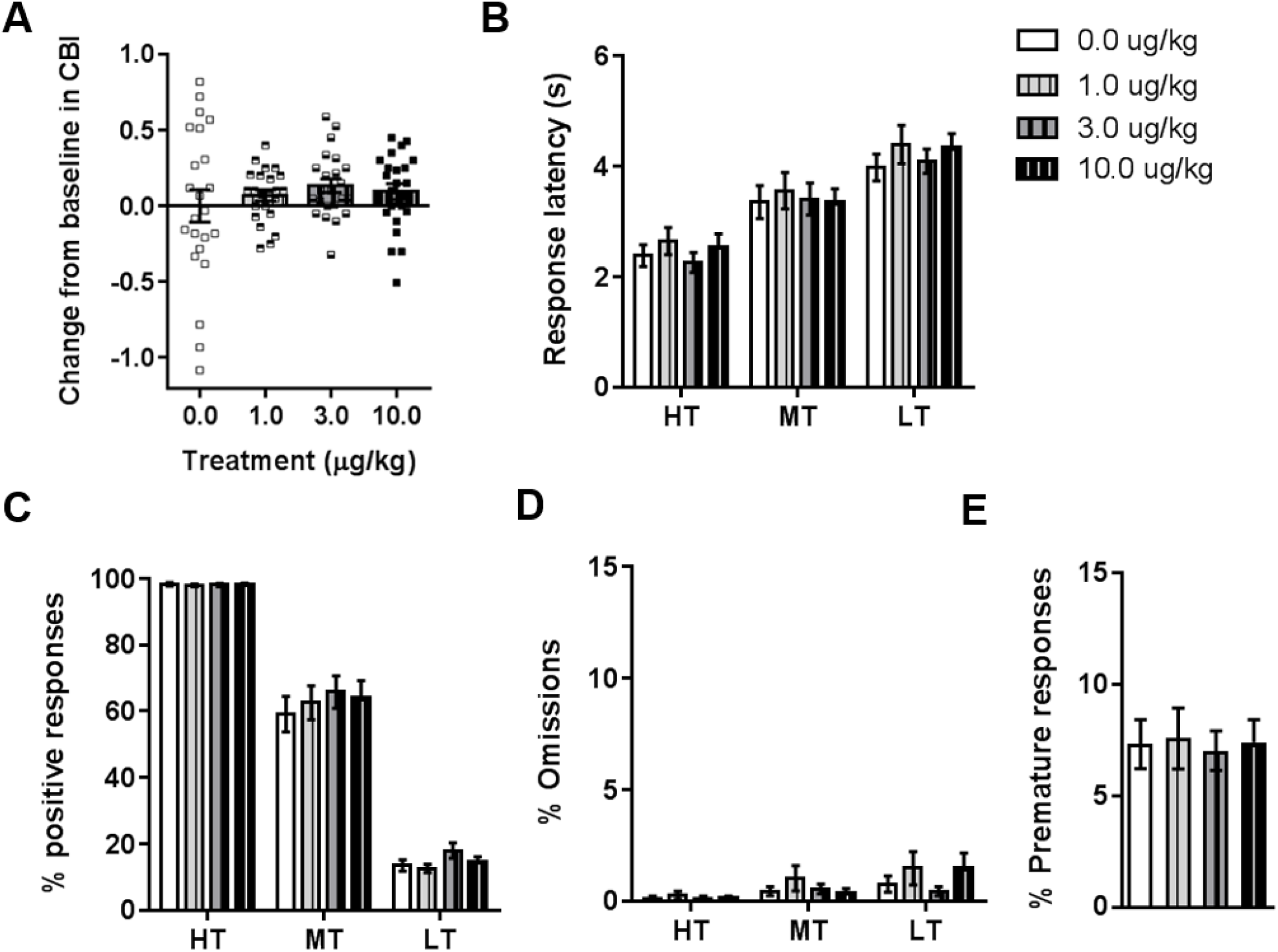
The effect of acute treatment with the immunomodulator lipopolysaccharide on judgement bias. Acute doses of lipopolysaccharide (LPS; 0.0, 1.0, 3.0, 10.0 μg/kg; n = 23) were administered by intraperitoneal injection prior to testing on the judgement bias task. (A-E) There were no effects on behavioural measures at any of the doses tested. Data shown and represent mean ± SEM (bars and error bars) overlaid with individual data points for each rat in panel A. 30 min pre-treatment. HT - high reward tone; MT - midpoint tone; LT - low reward tone.

**Figure 6.**
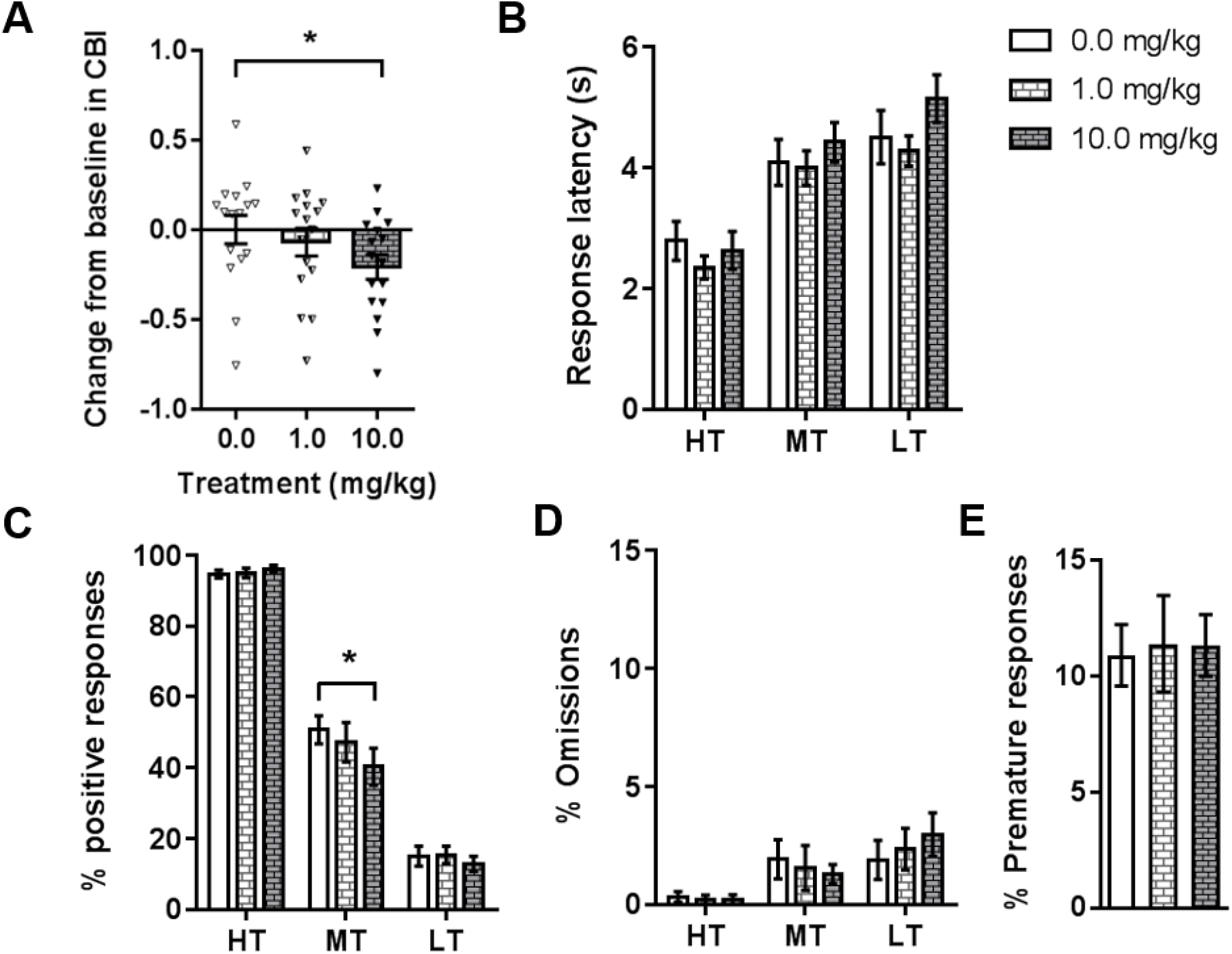
The effect of acute treatment with the immunomodulator corticosterone on judgement bias. Acute doses of corticosterone (CORT; 0.0, 1.0, 10.0 mg/kg; n = 16) were administered by subcutaneous injection prior to testing on the judgement bias task. (A) The 10.0 mg/kg dose of CORT caused a negative change in cognitive bias index (CBI) for the midpoint tone. (B) CORT did not alter response latencies. (C) CORT (10.0 mg/kg) reduced the percentage of positive responses for the midpoint tone (significant drug*tone interaction: *F*_4,60_=2.612, *p*=0.044 and post-hoc: *p*=0.026), which was also seen as in change in CBI (panel A). (D/E) CORT did not alter omissions or premature responses. Data shown and represent mean ± SEM (bars and error bars) overlaid with individual data points for each rat in panel A. 30 min pre-treatment. \**p*<0.05. HT - high reward tone; MT - midpoint tone; LT - low reward tone.

### Effect of chronic treatment with an immunomodulator on interpretation of the ambiguous cue in the JBT

Fifteen rats were initially split into control (n=8) and INF-α (n=7 groups). Data from one rat in the INF-α group could not be included as the animal died before the end of the study. Data from one other rat in the INF-α was excluded from the analysis as it did not meet accuracy criteria. These meant eight control animals and five INF-α animals were included in the final analysis.

There were no significant differences in any behavioural measures between groups in the pre-drug week (see Pre-drug sections of Figure 7 and Figure S6). There was a main effect of group (*F*_*1,11*_=5.297, *p*=0.042) and a trend towards a session*group interaction (*F*_*5,55*_ =2.077, *p*=0.082) across the entire study period (pre-drug, drug and post-drug) for change from baseline in CBI (Figure 7A). Analysing these data split by group revealed no effect of session for the control group (*F*_*1*.*860,13*.*022*_=0.324, *p*=0.714), but a main effect for the INF-α group (*F*_*5,35*_=3.579, *p*=0.018), indicating that CBI was changed across the experimental period in the INF-α treated rats. Post-hoc analysis showed that the INF-α group had a more negative change in CBI compared to control animals in the 3^rd^ and 4^th^ drug weeks, and in the post-drug week (independent samples t-tests: *p*=0.027, *p*=0.042 and *p*0.029 respectively). The INF-α treated animals were also more negative compared to their own baseline from drug week 2 onwards (one-sample t-tests: *p*s≤0.033). Chronic INF-α treatment did not cause changes in other behavioural measures, except for percentage positive responses for the midpoint tone (main effect of session: *F*_*5,55*_=5.379, p<0.001; and trend towards session*group interaction: *F*_*5,55*_=2.317, *p*=0.056; Figure 7C), which reflects the change in CBI. There were some changes in behavioural measures irrespective of treatment group. For all three tones, there was a main effect of session for response latency (*F*s≥2.799, *p*s≤0.025; Figure 7B and Figure S6A), indicating that both groups became quicker to respond over the entire study period. There was also a main effect of session for low tone omissions (*F*_*5,55*_=4.668, *p*=0.001), driven by both groups making fewer omissions in the drug and post-drug periods compared to the pre-drug week (Figure S6C).

**Figure 7.**
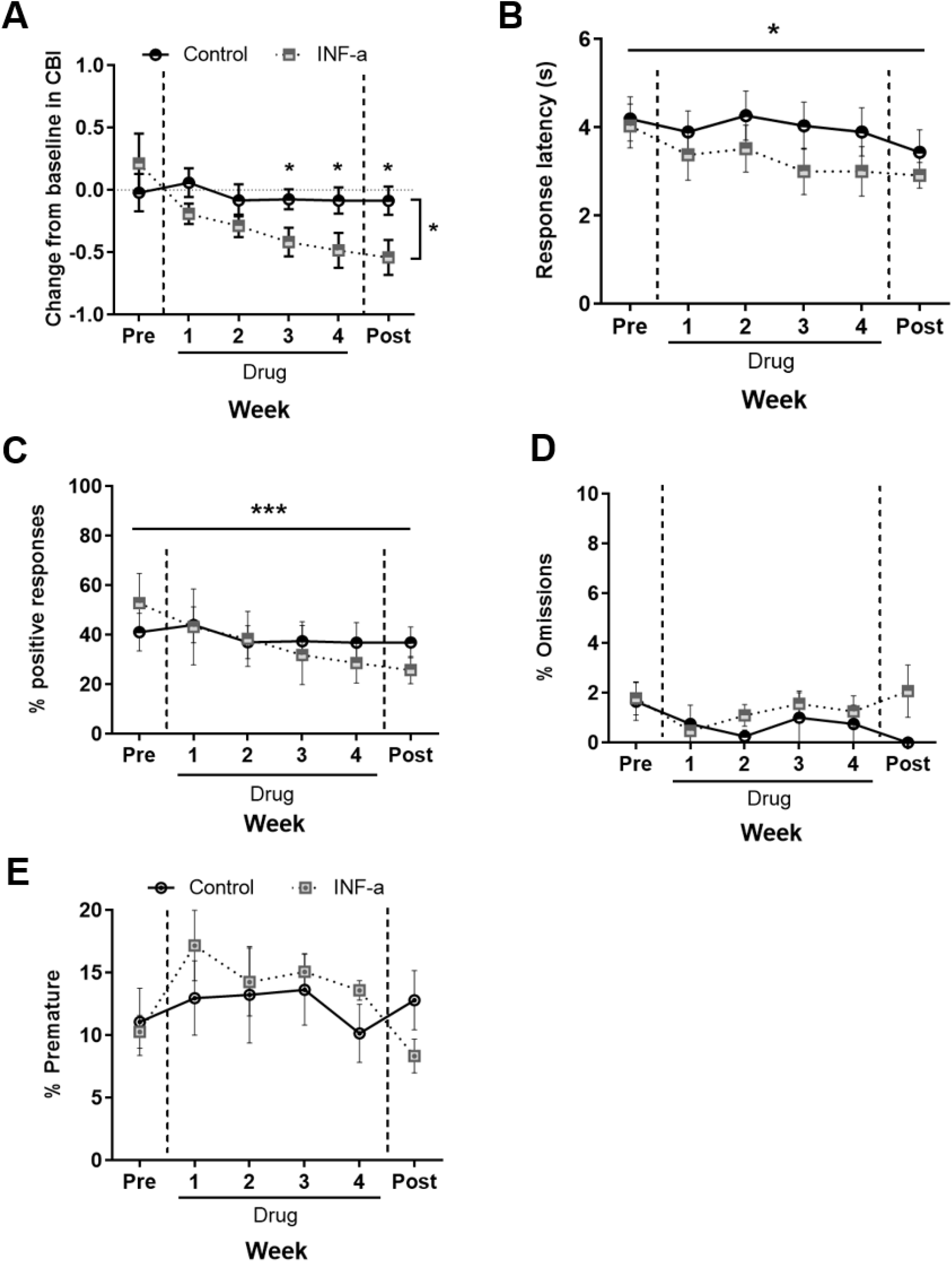
The effect of chronic treatment with the immunomodulator interferon-α on judgement bias. Rats assigned to the chronic interferon-α (INF-α) group experienced intraperitoneal injections of INF-α (100 units/kg) daily for four weeks, whilst control rats experienced daily intraperitoneal injections of saline vehicle (0.0 mg/kg). Twice weekly test sessions (averaged) were conducted one week prior to treatment (Pre), for the four weeks during treatment (Drug 1–4) and for one week following the end of treatment (Post). There were no significant differences between groups during the pre-drug period for any measure. (A) Rats in the INF-α group became more negative as treatment progressed (main effect of group: *F*_*1,11*_=5.297, *p*=0.042, trend towards session*group interaction: *F*_*5,55*=_2.077, *p*=0.082), with a more negative change in cognitive bias index (CBI) compared to controls during the 3^rd^ and 4^th^ drug week, and as well as during the post drug period (post-hoc comparisons: *p*=0.027, *p*=0.042, *p*=0.029 respectively). (B/C/D/E) Behavioural data for other measures are shown here for the midpoint tone only (see Supplementary Fig. 2 for data for reference tones). (B) Irrespective of treatment, rats became quicker to respond to the midpoint tone across weeks. (C) There was a main effect of session (*F*_*5,55*_ = 5.379, p < 0.001), and trend towards session*group interaction (*F*_*5,55*_ = 2.317, p = 0.056) for percentage of positive responses, reflecting the change in CBI shown in (A). (D) There were no differences in omissions between the control and INF-α groups. (F) There were also no changes in premature responding. Data shown are for the midpoint tone only, and represent mean ± SEM. Control group: n=8, INF-α group: n=5. ***p<0.001, *p<0.05.

## Discussion

Acute administration of pro-depressant drugs and immunomodulators that have previously been shown to induce learning and memory biases in the ABT^27,29^ failed to alter decision-making bias in the reward-based JBT, with the exception of acute treatment with corticosterone. Acute corticosterone treatment induced a negative judgement bias without altering other behavioural measures. Furthermore, chronic treatment with INF-α induced a negative decision-making bias in weeks 3 and 4 of treatment, and this negative bias lasted after treatment ended, despite acute treatment having no effect. This mirrors what has been seen previously in this JBT with psychosocial stress^19^ and conventional antidepressants^25^, where chronic but not acute manipulations altered judgement bias.

Acute treatments with pro-depressant drugs (rimonabant, retinoic acid and tetrabenazine) or immunomodulators (INF-α and LPS) did not induce change in decision-making bias in this JBT. This mirrors findings with acute restraint stress^19^, acute treatment with conventional antidepressants (fluoxetine, reboxetine and venlafaxine)^25^, and acute treatments with NMDA receptor antagonists (PCP, lanicemine, memantine)^26^ which all failed to change decision-making biases in this task. A potential common factor linking these results is the time course over which these drugs cause subjective reporting in change of mood or depression symptoms in humans. Patients only report improvements in depression ratings weeks to months after onset of treatment with conventional antidepressants^39^. Rimonabant was withdrawn as an anti-obesity drug after evidence that long-term treatment increased the risk of depressed mood disorders and anxiety^40^, and a later study showed that acute rimonabant treatment did not alter subjective reports of mood in humans^41^. Retinoic acid has been linked to an increased risk for depression, with most cases developing after 1-2 months of treatment^31^. For tetrabenazine, it has been reported that depression occurs in up to 15% of patients receiving long term treatment for Huntington’s disease^32,33^. IFN-α has been shown to induce depressive symptoms after weeks to months of treatment in 20–50% of patients^34^. LPS is not generally given to humans, instead being an endotoxin found on the cell wall of gram-negative bacteria, but acute treatment in rodents (at higher doses than used in this study) induces sickness-like behaviour^42^ that is thought to be comparable to bacterial infection in humans^43^. Chronic treatment with LPS has been used as a rodent model to induce depression-like behaviours^36^, and has been shown to cause reduced sucrose preference, a rodent test for anhedonia, following chronic, but not acute treatment^44^. The lack of effect on decision-making biases across all these drugs when given acutely, but the development of more negatively biased decision-making over a longer time period with chronic IFN-α treatment suggests that this reward-based JBT is sensitive to measuring affective biases that manifest on timescales that are more aligned with subjectively reported mood change in humans.

These findings contrast effects on learning and memory biases seen in the ABT, where acute treatment with the conventional antidepressants above did induce positive affective biases^27^, whilst the same pro-depressant drugs and immunomodulators (tested at the same doses) induced negative affective biases^27,29^, suggesting these two tasks are measuring distinct types of affective bias. Evidence from these two tasks suggests that learning can be modified acutely whilst alterations in decision-making take longer. This could be due to the nature of the two tasks: in the JBT, decision-making about the ambiguous cue requires the animal to have learnt, over a long training period, outcomes about two other related cues, and recall this information to make a judgement about their choice on an ambiguous trial. However, in the ABT, specific memories (one following treatment, one following control) that have been learnt over only four sessions (two per manipulation) are being tested^27^. These are likely to be modifiable on a much shorter time scale. The implications relating to treatment of MDD are important, as both types of bias are likely to play a role in the pathology of the disorder, but decision-making biases may be slower to change, whilst potentially also being more likely to impact on people’s behaviour. For example, if a person subjectively feels more negative, and hence pessimistic, they will be less likely to make good decisions.

The clear-cut, but contrasting effects of these drug treatments on two tasks measuring affective biases in rats are not reflected in other rodent tasks traditionally used to measure depression-like behaviour, such as the forced swim test (FST), a measure of behavioural despair, and the sucrose preference test (SPT), a measure of anhedonia. For example, in preclinical studies acute rimonabant has been shown to either have no effect on depression-like behaviours, or even to show antidepressant-like effects (e.g. Griebel et al.^45^ and references within). However, chronic rimonabant treatment reduced sucrose consumption and increased immobility time in the FST^46^. For IFN-α treatment, there are reports showing that acute and chronic treatment regimes in rodents increase immobility in the FST^47-49^ and reduce sucrose preference^49^, as well as reports that show no effect^51^. This adds to growing evidence that traditionally used preclinical tests such as the FST and SPT may not be the most reliable and lack aspects of validity for detecting pro-depressant or antidepressant efficacy across drugs with a range of pharmacological actions^52^. Instead, tasks measuring affective biases, which are translatable across species, may be more useful measures in preclinical research.

The exception to the lack of effects on decision-making biases with pro-depressant drugs and immunomodulators was acute treatment with corticosterone, which did induce a negative bias. Acute, negative decision-making biases have also been seen in this reward-based JBT with FG7142^19^, an anxiogenic drug that acts as a partial inverse agonist at the GABA_A_ receptor, as well as acute treatments with noradrenergic drugs in the reward-punishment JBT, including reboextine^53^, desipramine^54^, and co-treatment with reboxetine and corticosterone^55^. This finding therefore partially replicates previous findings by Enkel et al.^55^, but also suggests that negative decision-making biases can be induced by direct activation of the stress system.

## Conclusions

Overall, these findings back up previous studies using the reward-based JBT that have shown differential effects of acute and chronic pharmacological manipulations on decision-making biases.

## Acknowledgements

This research was funded by an Industrial Partnership Award awarded by BBSRC in collaboration with Boehringer Ingelheim (Grant no: BB/N015762/1) and carried out with intellectual support from Boehringer Ingelheim.

## Data Availability Statement

Supporting data available on request.

## Author contributions statement

CAH ran the experiments, analysed data, and wrote and edited the manuscript. JMB ran the experiments and analysed data. RA and BH formulated the overall concept and read the paper. ESJR formulated the overall concept, and wrote and edited the paper.

