## Supplementary Figures and Tables for "Effects of pro-depressant and immunomodulatory drugs on biases in decision-making in the rat judgement bias task"

**Supplementary Table S1**

| Stage | Description | Criteria | Sessions required to meet criteria | | |
|---|---|---|---|---|---|
| | | | Cohort 1 | Cohort 2 | Cohort 3 |
| 1 – Magazine training | Tone (2 kHz only for half the session followed by 8 kHz only for the rest of the session, order counterbalanced across rats) played for 20 s followed by release of one pellet into magazine; 10 s ITI. No levers available. | 20 pellets eaten for each tone frequency | 1 | 1 | 1 |
| 2 – Tone training | Response on lever during tone (2 kHz or 8kHz only, order counterbalanced across rats) rewarded with one pellet. Lever corresponding to that tone frequency available only. | > 50 trials completed for two consecutive sessions on each tone frequency | 4 | 4 | 4 |
| 3 – Discrim-ination training | Response on correct corresponding lever only during tone (either 2 kHz or 8 kHz presented pseudorandomly) rewarded with one pellet. Both levers available. Incorrect or omitted trials were repeated (i.e. same tone frequency played) until a correct response occurred. | > 70% accuracy for both tones, no significant differences on analysed behavioural measures over three sessions and < 1:1 ratio of correct:premature responses | 15 | 15 | 10 |
| 4 – Reward magnitude training | As Stage 3 but response on correct corresponding lever only rewarded with four pellets for high reward tone and one pellet for low reward tone. Both levers available. | As for Stage 3 but with > 60% accuracy for both tones (to allow for biases in responding to reference tones caused by the difference in associated reward magnitude). | 9 | 9 | 10 |
| 5 – Baseline session | Same format as reward magnitude training sessions. | Animals had to show equivalent baseline session performance to pre-drug study baseline sessions (measured by no significant differences pre- and post- on behavioural measures. | - | - | - |
| 6 – Probe sessions | For reference tones (2 of 8 kHz) response on correct corresponding lever during the tone rewarded with either 4 pellets (2 kHz) or 1 pellet (8 kHz) reward. For ambiguous midpoint tone (5 kHz), random reinforcement was used whereby outcomes for 50% of the trials followed 2 kHz tone trials, whilst the other 50% followed 8 kHz tone trials (see Supplementary Figure XX for further details). There were no repeated trials following incorrect or omissions. | < 60% accuracy for both reference tones, < 50% omissions and | - | - | - |

For all training stages, trial structure was as depicted in Figure S1 (except for magazine training which excludes any form of lever press response). Pseudorandom tone presentation was achieved by splitting each session into blocks of 10 trials, within which there were 5 presentations of each tone frequency (2 or 8 kHz). Within a block, there could only be a maximum of n-1 consecutive tone presentations (i.e. in this trial structure, a maximum of 4 consecutive trials of either 2 or 8 kHz). Omitted trials (no lever press during tone presentation; possible in stages 2-4) were punished with a 10 second timeout where the house light was turned on, and the animal was unable to initiate another trial. Incorrect trials (wrong lever for the tone presented; possible in stages 3-4) were also punished with a 10 second timeout with the house light on. Premature trials (lever press during the ITI; possible during stage 2-4) were similarly punished with a 10 second timeout with the house light on. Each new trial had to be self-initiated by the animal by making an entry into the magazine (this was signalled by the magazine light being turned on, and was switched off once animals made the magazine nose poke). Baseline session were the same format as reward magnitude training sessions, with animals required to repeat incorrect or omitted trials. Training stages 1-5 consisted of a maximum of 100 trials, or 60 minutes, and where both tones were played (stages 3-5) tone type was equally spilt across the session (50 trials per tone). Probe sessions consisted of 120 trials: 40 of each reference tone (2 and 8 kHz) and 40 midpoint tones (5 kHz).

**Supplementary Table S2**

| Behavioural measure | Analysed for: | Description | Statistical analysis |
|---|---|---|---|
| **Response latency** | Each tone | Time between presentation of the tone and response on the lever (correct lever for high and low reward tones, either lever for midpoint tone) | Two-way repeated measures ANOVA with tone and session as within-subjects factors |
| **Percentage positive responses** | Each tone | Number of responses made on the high reward lever divided by the total number of responses made and on the high reward and low reward levers for that tone | |
| **Percentage omissions** | Each tone | Number of trials where no lever press occurred during 20 s tone presentation divided by total completed trials for that tone | |
| **Percentage of premature responses** | Whole session | Number of trials where a response was made in the 5 s inter-trial interval divided by total completed trials | Repeated measures ANOVA with session as the within-subjects factor (or paired samples t-test if only two drug treatments) |

This table details the other behavioural measures (apart from cognitive bias index) that were analysed for each experimental manipulation. ANOVA – analysis of variance.

**Supplementary Figure S1**

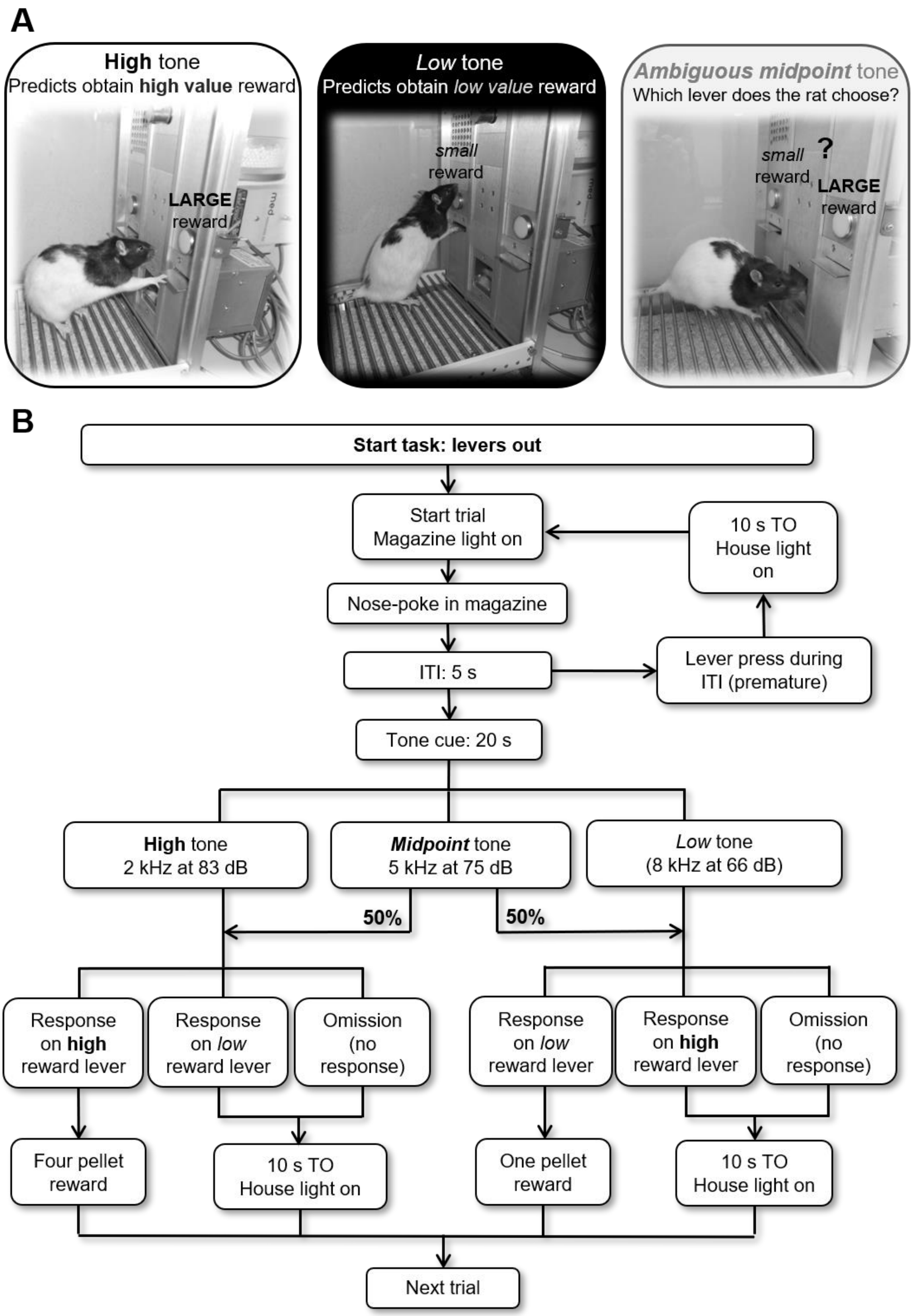


**Supplementary Figure S1 – Schematic of the JBT and trial structure.**

In the judgement bias task (JBT), rats are trained to associate one tone frequency (2 kHz) with a high value reward: i.e. if the rat presses the correct lever (shown as the left lever in (A), but counterbalanced across rats in a cohort) they receive a high value reward (four reward pellets). They also learn to associate a second tone frequency (8 kHz) with receiving a low value reward (one reward pellet; shown in (A) as pressing the right lever during the tone). Judgement bias, or decision making about an ambiguous cue, which is known to be influenced by affective state, can be probed by presenting an ambiguous tone that has a midpoint frequency between the two reference cues (5 kHz), and recording which lever the rat presses. If the rat is expecting the more positive outcome (indicative of an optimistic judgement bias), then they will more often choose the large reward lever, but if the rat is in a more negative affective state, they will expect the less positive outcome and more often choose the low reward lever, a pessimistic judgement bias. During the task, tones are presented within discrete trials, the format of which is depicted as a flow chart in (B). The task is self-initiated, and so each trial begins only once the rat makes a nosepoke entry into the magazine port. This is followed by a 5 second inter-trial interval (ITI), during which time the rat has to wait and refrain from making a lever press response. If the rat does press a lever, they are punished with a 10 second timeout (TO). The tone cue is presented for a maximum of 20 seconds following the ITI, or until the rat makes a lever press response. The outcome following each lever press depends on which tone was played, and which lever was pressed. Correct lever presses to either reference tone (high or low tones) results in the corresponding reward being delivered to the magazine, whilst incorrect lever presses results in a 10 second TO. This TO also occurs if the rat fails to make any lever press during the 20 second tone presentation (an omission). During TOs, lever presses and magazine entries are recorded but have no consequences, meaning the rat has to wait to be able to begin the next trial. When the midpoint tone is presented, 50% of the time this tone is “classified” by the software as having the same response properties as the high reward tone. I.e., if the rat makes a high reward lever press during a midpoint tone presentation classified in this way, then they will receive a four pellet reward, but will experience the 10 second TO if they make a low reward lever press. Similarly, if the midpoint tone is “classified” as having the same response properties as the low reward tone, then a high reward lever press would result in a TO, whilst a low reward lever press would result in delivery of the small reward. In this way, each lever is only ever associated with the same reward outcome (i.e. four pellets for the high reward lever), but the midpoint tone becomes randomly reinforced, so rats will maintain responding for this tone across multiple trials within a session, whilst being unable to learn a specific reward contingency to associate with the midpoint tone. Image from Hales et al., 2020[26].

**Supplementary Figure S2**

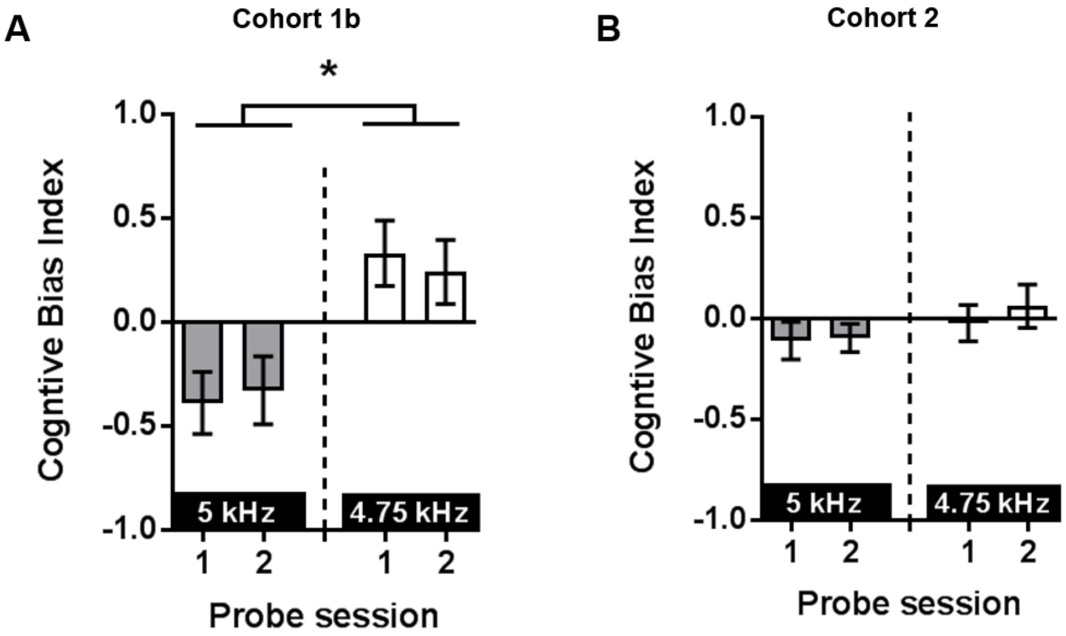


**Supplementary Figure S2** – *Comparison of performance on judgement bias probe tests with no experimental manipulation with different frequency midpoint tones.*

The midpoint ambiguous tone frequency was altered from 5 kHz to 4.75 kHz to adjust for the unusually negative baseline cognitive bias index (CBI) of cohort 1b. Data is shown for two probe tests conducted without any experimental manipulation for each frequency of midpoint tone for both this cohort, and for cohort 2 (which were adjusted to make data consistent for experimental manipulations that used both of these cohorts together). (A) Altering the frequency of the midpoint tone caused CBI to become more positive for cohort 1b (main effect of frequency: $F_{1,7}$=10.297, $p$=0.015). (B) Altering the midpoint tone frequency did not have any significant effect on cohort 2. Data shown and represent mean ± SEM. *$p$<0.05. HT - high reward tone; MT - midpoint tone; LT - low reward tone.

**Supplementary Figure S3**

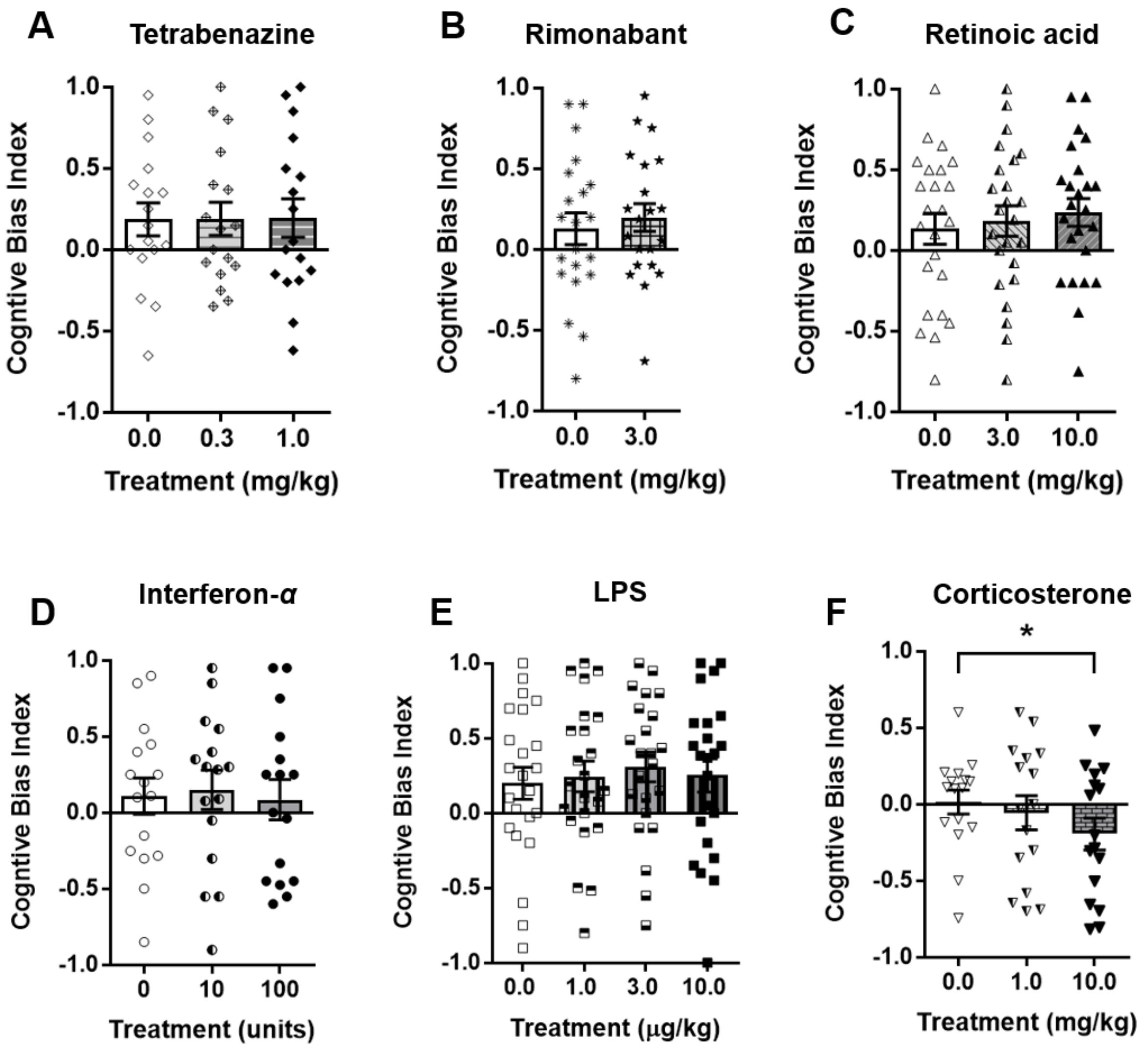


**Supplementary Figure S3** – *Raw CBI data for all acute drug studies in the JBT.*

Acute doses of (A) tetrabenazine (0.0, 0.3, 1.0 mg/kg), (B) rimonabant (0.0, 3.0 mg/kg), (C) retinoic acid (0.0, 3.0, 10.0 mg/kg), (D) interferon-*α* (INF-*α;* 0, 10, 100 units/kg), (E) lipopolysaccharide (LPS; 0.0, 1.0, 3.0, 10.0 ug/kg), or (F) corticosterone (CORT; 0.0, 1.0, 10.0 mg/kg; n = 16) were administered by intraperitoneal (A-E) or subcutaneous injection (F) prior to testing on the judgement bias task. This data is the raw cognitive bias index (CBI) data from Figures 1-6. (A-E) None of the drugs tested CBI to be altered, except for (F) acute treatment with 10.0 mg/kg CORT. Data shown and represent mean ± SEM (bars and error bars) with individual data points overlaid. *$p$<0.05.

**Supplementary Figure S4**

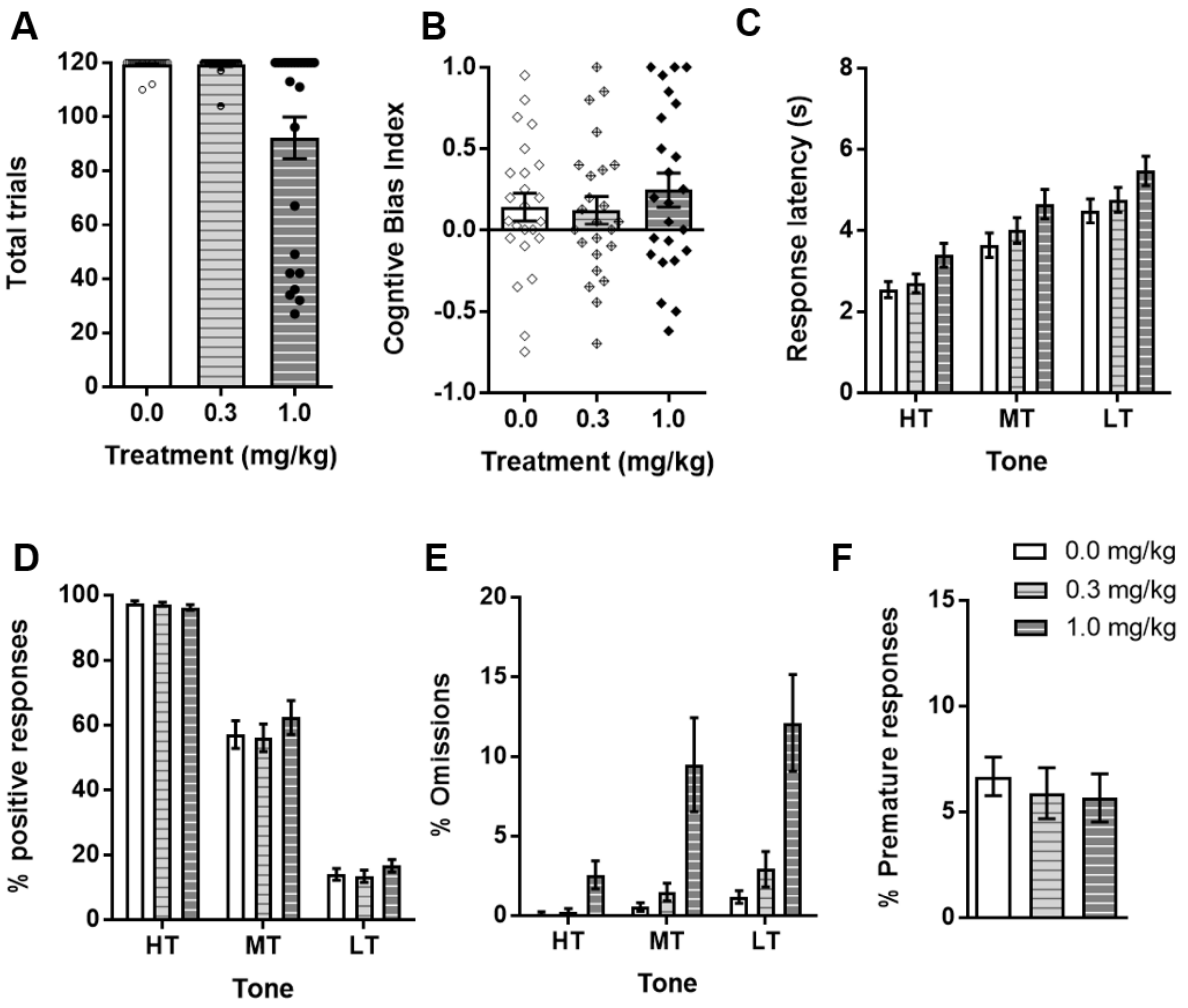


**Supplementary Figure S4** – *Full dataset without exclusions for the effect of acute treatment with the putative pro-depressant drug tetrabenazine on judgement bias.*

Acute doses of tetrabenazine (0.0, 0.3, 1.0 mg/kg; n = 24) were administered by intraperitoneal injection prior to testing on the judgement bias task. (A) The highest dose (1.0 mg/kg) caused some animals (n = 7) to perform less than half of the total trials, hence them being excluded from the main analysis. (B-F) Behavioural measures without exclusions for the tetrabenazine drug study: (B) cognitive bias index (CBI); (C) latency to respond; (D) percentage of positive responses made to each tone; (E) percentage omissions and (F) percentage of premature responses. The high dose (1.0 mg/kg) also caused a large increase in missed trials (E). Data shown and represent mean ± SEM (bars and error bars) overlaid with individual data points for each rat on panels A-B. 30 min pre-treatment. HT - high reward tone; MT - midpoint tone; LT - low reward tone.

**Supplementary Figure S5**

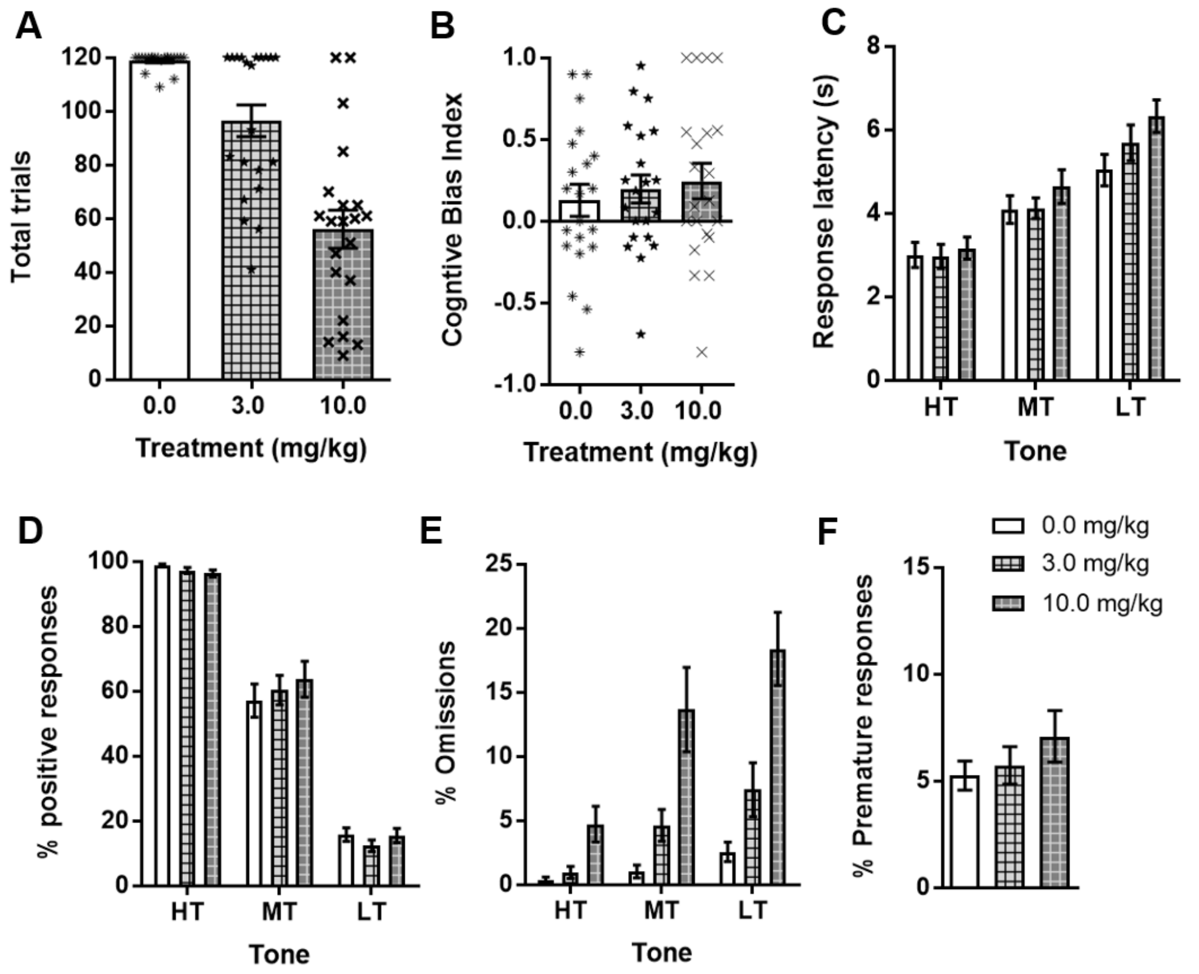


**Supplementary Figure S5** – *Full dataset without exclusions for the effect of acute treatment with the putative pro-depressant drug rimonabant on judgement bias.*

Acute doses of rimonabant (0.0, 3.0, 10.0 mg/kg; n = 24) were administered by intraperitoneal injection prior to testing on the judgement bias task. (A) The highest dose (10.0 mg/kg) caused more than half of animals to perform less than half of the total trials, hence the highest dose was excluded from the main analysis. (B-F) Behavioural measures without exclusions for the rimonabant drug study: (B) cognitive bias index (CBI); (C) latency to respond; (D) percentage of positive responses made to each tone; (E) percentage omissions and (F) percentage of premature responses. The highest dose (10.0 mg/kg) also caused a large increase in missed trials (E). Data shown and represent mean ± SEM (bars and error bars) overlaid with individual data points for each rat on panels A-B. 30 min pre-treatment. HT - high reward tone; MT - midpoint tone; LT - low reward tone.

**Supplementary Figure S6**

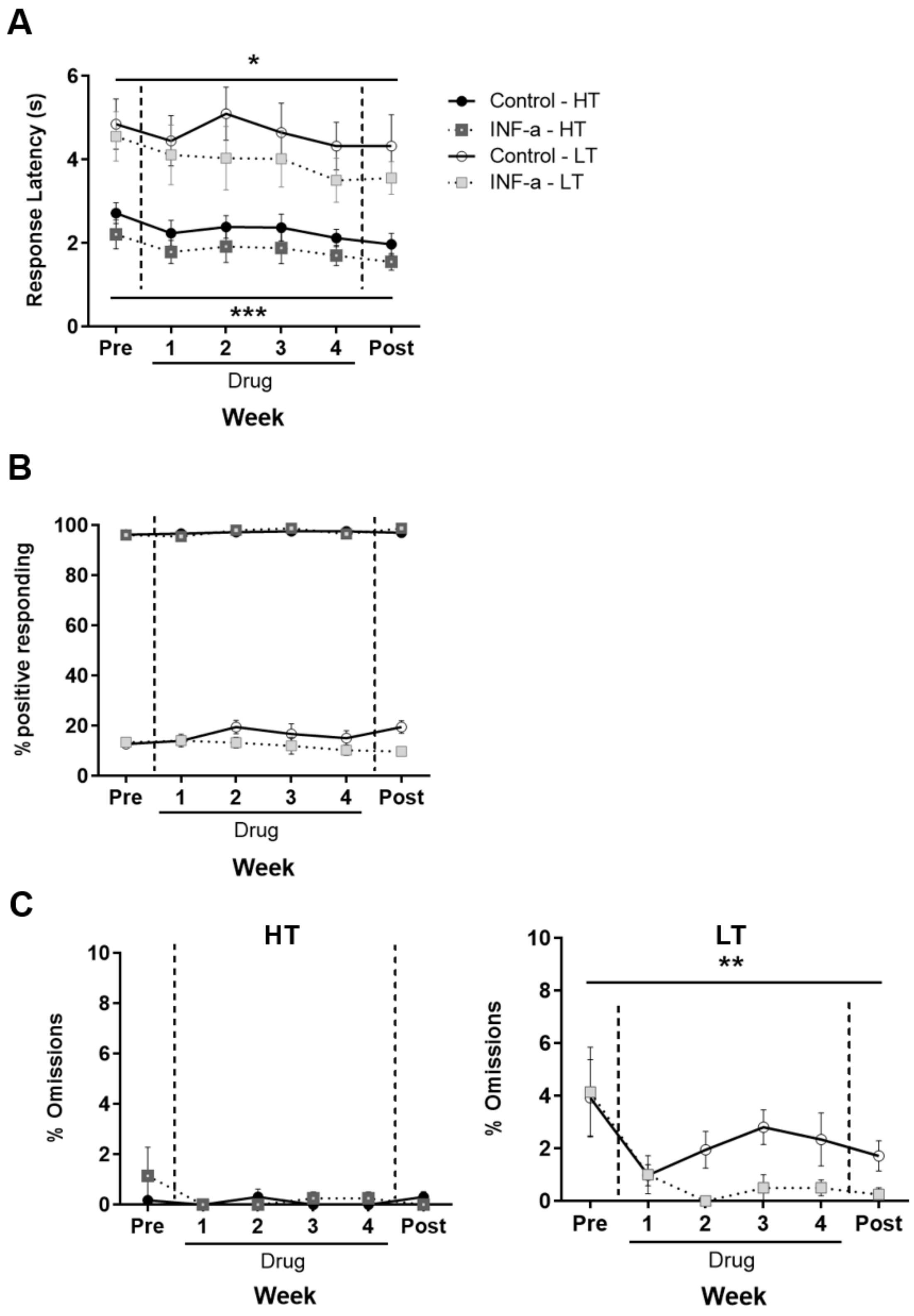


**Supplementary Figure S6** – *Behavioural measures for reference tones for chronic treatment with interferon-α.*

Rats assigned to the chronic interferon-α (INF-α) group experienced intraperitoneal injections of INF-α (100 units/kg) daily for three weeks, whilst control rats experienced daily intraperitoneal injections of saline vehicle (0.0 mg/kg). Twice weekly test sessions (averaged) were conducted one week prior to treatment (Pre), for the four weeks during treatment (Drug 1–4) and for one following the end of treatment (Post). There were no significant differences between groups during the pre-drug period for any measure. (A) There was a main effect of response latency for both reference tones(high: $F_{5,55}$=5.365, $p$<0.001; low: $F_{5,55}$=2.799, $p$=0.025), showing that irrespective of treatment, rats became quicker to respond across weeks. (B) There were no changes in percentage of responses made for the reference tones. (C) There was no difference in omissions for the high reward tone (left graph), but a main effect of session ($F_{5,55}$=4.668, $p$=0.001) indicated that across both groups, rats made fewer omissions for the low tone across weeks (right graph). Data shown represent mean ± SEM. **$p$<0.01, *$p$<0.05. HT - high reward tone; LT - low reward tone.
